# T helper 2 transcriptional profile predicts single-cell HIV envelope-specific polyfunctional CD4+ T cells correlated with reduced risk of infection in RV144 trial

**DOI:** 10.1101/2021.06.07.447395

**Authors:** Kristen W. Cohen, Yuan Tian, Casey Thayer, Aaron Seese, Robert Amezquita, M. Juliana McElrath, Stephen C. De Rosa, Raphael Gottardo

## Abstract

Despite the critical role antigen-specific T cells play in responding to viral infections, their aggregate frequencies in peripheral blood have not correlated with clinical protection during HIV infection. However, a subset of HIV-specific CD4+ T cells, termed polyfunctional T cells, can produce multiple effector cytokines simultaneously. In the RV144 HIV vaccine trial, polyfunctional T cells correlated with reduced risk of HIV infection. Little is known about what differentiates polyfunctional T cells from other vaccine-elicited T cells. Therefore, we developed a novel live-cell multiplexed cytokine capture assay, to identify and transcriptionally profile vaccine-specific polyfunctional CD4+ T cells. We applied these methods to samples from the HVTN 097 clinical trial of the same vaccine regimen as RV144. We discovered two surface receptors that were enriched among polyfunctional CD4+ T cells and a Th2-biased signature (*IL-4, IL-5*, and *IL-13*) that specifically predicted the envelope-specific polyfunctional CD4+ T cells that were correlated with reduced risk of HIV infection in RV144. By linking single-cell transcriptional and functional profiles, we may be able to further define the role of vaccine-elicited polyfunctional T cells in contributing to effective immunity.

**Key Points:**

- Novel *ex vivo* multiplexed cytokine capture assay to enumerate and single-cell sort polyfunctional T cells for downstream analyses
- Polyfunctional T cells were specifically detected among the HIV envelope-stimulated CD4+ T cells
- Single-cell RNA sequencing identified novel surface markers enriched among vaccine-specific polyfunctional CD4+ T cells
- Th2 transcriptional signature predicted polyfunctional CD4+ T cell profile that had correlated with reduced risk of HIV infection in the RV144 HIV efficacy trial

## Introduction

Polyfunctional HIV-specific T cells are able to simultaneously produce multiple functions or cytokines and have been repeatedly correlated with improved clinical outcomes in HIV infection (1, 2) and vaccination (3). Utilizing an unbiased, Bayesian statistical approach, referred to as COMPASS, we previously determined that polyfunctional HIV envelope (Env)-specific CD4+ T-cell responses correlated with a reduced risk of HIV infection in the RV144 trial (4), which is the only HIV vaccine trial, thus far, that has demonstrated vaccine-mediated efficacy, although modest (5). In particular, Env-specific CD4+ T cells with two highly polyfunctional cytokine profiles correlated with reduced risk of HIV infection: 1) IFN-γ, TNF-α, IL-4, IL-2, and CD154 [odds ratio (OR): 0.57, p=0.006] and, 2) IL-4, IL-2, and CD154 (OR: 0.63, p=0.013) (4). To gain a deeper understanding of the mechanism(s) by which Env-specific polyfunctional CD4+ T cells may be contributing to protective immune responses to HIV vaccination, we were interested in performing single-cell transcriptional analyses of vaccine-induced polyfunctional CD4+ T cells. There may be signatures that lie within the transcripts of these polyfunctional cells that are vital to vaccine efficacy compared to cells expressing one or two effector functions. However, the intracellular cytokine staining (ICS) methods used to identify these cell subsets require fixation and permeabilization, which damages the mRNA and precludes downstream transcriptional analysis. Alternatively, activation induced marker (AIM) assays have been used to identify and sort antigen-specific T cells for transcriptional analyses (6-8). These assays rely on upregulation of surface markers that are preferentially upregulated after activation. However, these markers have variable baseline expression which effects the specificity of detection of vaccine-specific cells and require extended incubation times (up to 18 hours), which can alter the transcriptional profiles as well as result in bystander activation of cells that are not vaccine-specific. Lastly, these methods do not allow for polyfunctional T cells to be specifically identified and thus interrogated.

Therefore, we developed a novel multiplexed cytokine capture assay to be able to identify and sort individual, viable, antigen-specific polyfunctional CD4+ T cells for single-cell RNA sequencing (scRNA Seq). In addition, the single cells were index-sorted, enabling scRNA sequence data to be linked to the protein expression observed in the flow cytometry results. The resulting analyses demonstrated that polyfunctional T cells could be specifically detected among the vaccine-stimulated T cells by the multiplexed cytokine capture assay. It also led to the identification of novel surface-expressed activation markers, CD82 and CD44, which were significantly enriched among the polyfunctional CD4+ T cells. Lastly, we established a Th2 transcriptional signature that predicted the polyfunctional CD4+ T cell profiles that had correlated with reduced risk of HIV infection in the RV144 HIV efficacy trial.

## Results

### Evaluation of vaccine-specific CD4+ T cells by multiplexed cytokine capture assay

HVTN 097 was a phase 1b clinical trial that tested the prime-boost vaccine regimen that was used in the RV144 efficacy trial: ALVAC-gp140 boosted with the co-administered bivalent gp120 proteins, as described previously (5, 9). One of the primary goals of HVTN 097 was to confirm the immune profiles induced by this vaccine regimen in RV144, originally conducted in Thailand, in a different recipient population, South Africa. Indeed, the vaccine regimen in HVTN 097 induced Env-specific polyfunctional CD4+ T cell responses, including the specific subsets which had correlated with reduced risk of infection in RV144 (9, 10).

To interrogate the transcriptional profiles of these rare subsets of cells, we developed a live-cell multiplexed cytokine capture assay to detect and sort vaccine-specific polyfunctional CD4+ T cells (Figure 1). To detect CD154 (CD40L), we preincubated the cells with antibody against CD40 to block uptake of its ligand, CD154, and then co-cultured the cells with fluorescently-labeled anti-CD154 antibody during the stimulation. Then, we multiplexed cytokine capture reagents to simultaneously detect secreted IFN-γ, IL-2 and IL-4. Due to experimental limitations, we decided to not use TNF-α as it was commonly co-expressed with IFN-γ among the polyfunctional T cell subsets and was deemed not critical for distinguishing the Env-specific CD4+ T cell “correlate” profiles: IFN-γ, TNF-α, IL-4, IL-2, and CD154 (OR: 0.57, p=0.006) and IL-4, IL-2, and CD154 (OR: 0.63, p=0.013) (4). Using this multiplexed cytokine capture assay, we single-cell sorted live cytokine-positive Env-specific CD4+ T cells from a subset of HVTN 097 vaccine recipients, 2 weeks post-final vaccination, for transcriptional profiling by scRNA Seq (Figure 1). The sample subset was selected based on previous ICS data showing the presence of vaccine-induced Env-specific CD4+ T cells and sample availability post-vaccination.

**Figure 1.**
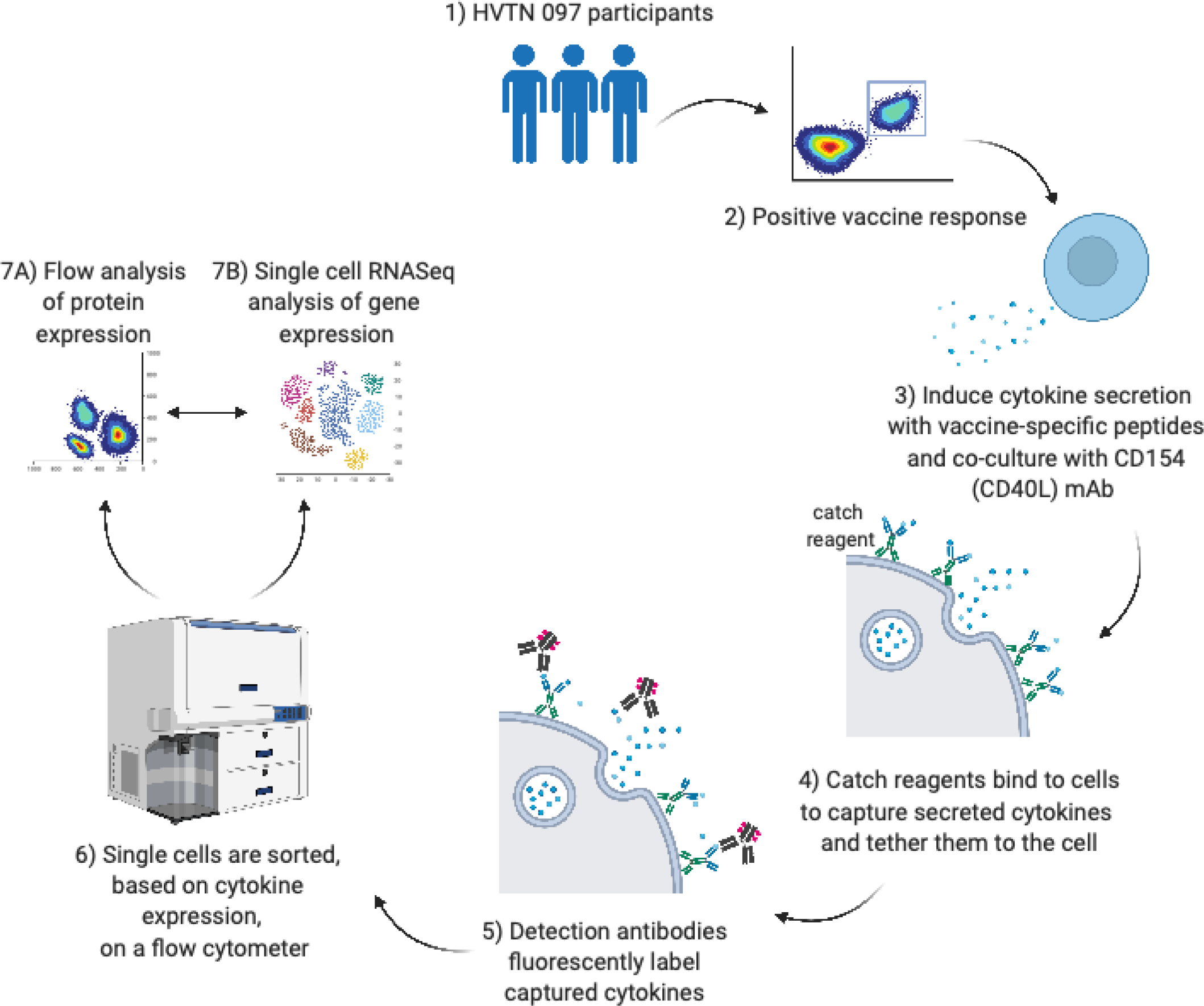
Overview of multiplexed cytokine capture assay and scRNA-Seq approach. Participants were enrolled in the HVTN 097 clinical trial testing the RV144 vaccine regimen (1) and samples were selected for having HIV Env-specific CD4 T cells by ICS assay (2). The multiplex cytokine capture assay uses antigen-specific peptides to induce the secretion of cytokines by vaccine-specific T cells (3). The multiplexed catch reagents adhere to the cells and capture the secreted IL-2, IL-4 and IFN-*γ* cytokines to the cell surface (4). The captured cytokines are then detected by fluorescently-labeled antibodies (5). Cytokine-positive cells are single-cell sorted by FACS (6) and sequenced by scRNA-Seq (7).

Initially, we characterized the cytokine profiles of the Env-specific CD4+ T cells detected using the multiplex capture assay by flow cytometry. In general, we were able to detect diverse functional Env-specific CD4+ T cell responses (Figure 2A and 2B). While there was some background detected in the monofunctional individual cytokines, the background was low or non-existent among polyfunctional subsets (Figure 2A). The magnitudes of Env-specific CD154+ and IL-4+ CD4+ T cells were similar between the two assays (p=ns), whereas frequencies of IL-2+ and IFN-γ+ CD4+ T cells were higher in the ICS assay compared to the capture assay (p=0.03; Figure 2B). The higher IL-2+ and IFN-γ+ frequencies in the ICS assay were likely due to the longer time during which secretory inhibitors allow cytokines to accumulate intracellularly for detection in the ICS assay (6 hours). In contrast, in the multiplexed cytokine capture assay, the cytokine capture reagents are only added for the last hour of the incubation and are also limited by the number of the catch reagents bound to the surface of the cell. CD154 is a surface-expressed T cell activation marker which can be detected by intracellular staining as is done in the ICS assay. In the multiplexed cytokine capture assay, we used co-culture with a fluorophore-labeled anti-CD154 detection antibody. Detection of CD154 by the two methods yielded similar frequencies and represented the single largest population of responding CD4+ T cells in both assays (Figure 2B and 2C). In contrast, there were more monofunctional IFN-*γ*+, IL-2+ and IL-4+ T cells detected in the multiplexed cytokine capture assay compared to ICS (Figure 2C and 2D). In general, there was a similar distribution of polyfunctional profiles among Env-specific T cells in the capture assay compared to the ICS assay (Figure 2D). Indeed, the multiplexed cytokine capture assay had reliable detection of diverse polyfunctional vaccine-specific CD4+ T cells in each of the samples tested, including the two highly polyfunctional (3 and 4-function) subsets which had correlated with reduced risk of infection in RV144: IFN-γ, IL-4, IL-2, and CD154; and IL-4, IL-2, and CD154 (Figure 2D). Lastly, the multiplexed cytokine capture assay had negligible CD4+ T cells expressing 2 or more cytokines in the negative control stimulations indicating the highly specific detection of polyfunctional T cells in the Env-stimulated samples (Figure 2A).

**Figure 2.**
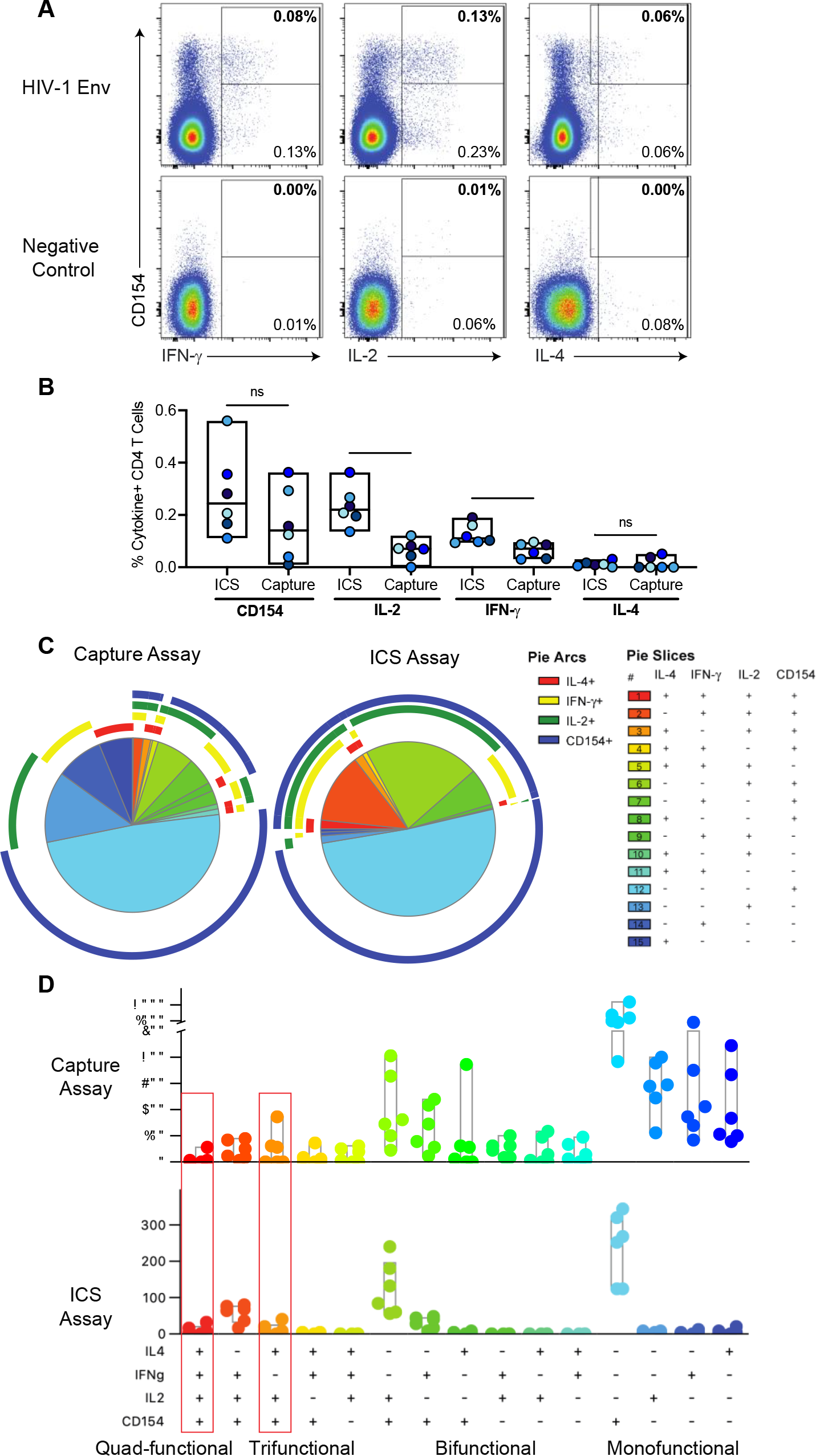
Comparison of Env-specific CD4+ T cell responses post-vaccination by multiplexed cytokine capture and ICS assays. A) Example of multiplexed cytokine capture assay Env and negative control stimulations of PBMC from a HVTN 097 vaccine recipient. Data is gated on CD4+ T lymphocytes (live single, CD56-, CD19-, CD14-, CD3+, CD4+ and CD8-). The Y-axis shows CD154 expression, and the X-axis shows: IFN-γ (left), IL-2 (middle), and IL-4 (right). The percentage in the bottom right of each plot indicates the frequency of cytokine positive events and the percentage in the top right (in bold) of each plot is the percentage of CD4 T cells that are co-expressing each cytokine with CD154. B) Background-subtracted percent of CD4 T cells expressing CD154, IFN-γ, IL-2, and IL-4 for each subject by multiplexed cytokine capture and ICS assays (n=6). Significance determined using Wilcoxon Rank test. * indicates p value < 0.05. C) Pie charts show the median distribution of subsets among the Env-specific CD4 T cells. Each pie slice indicates the proportion of Env-specific CD4 T cells that expressing those cytokines (indicated in the legend) and color-coded by the degree of polyfunctionality: monofunctional (blue), bi-functional (green), tri-functional (yellow/orange) and quad-functional (red). The pie arc colors allow visualization of the functions expressed by each slice. D) Bar chart of the per subject counts of Env-specific CD4+ T cell subsets by functional profile from ICS or multiplexed cytokine capture assay and color-coded by the degree of polyfunctionality. The red boxes indicate the two subsets that correlated with reduced risk of HIV infection in the RV144 vaccine trial.

### Single-cell gene expression analysis of vaccine-induced CD4+ T cells with diverse functional profiles

We used a combination of sort criteria in the multiplexed capture assay to single-cell index-sort a diverse representation of cytokine-positive Env-stimulated CD4+ T cells for transcriptional profiling by scRNA Seq. Since the polyfunctional CD4+ T cells are very rare, after sorting we preferentially selected polyfunctional T cells in addition to a random selection of other cytokine-positive T cells for conducting scRNA Seq. We conducted reverse transcription using SMART primers which incorporate a universal primer sequence to ensure sensitive and unbiased cDNA synthesis and subsequently amplified the whole transcriptome (11).

We performed scRNA Seq on Env-specific CD4+ T cells with diverse functional profiles from the post-vaccination samples. The distribution of Env-specific CD4+ T cells (n=1,423) for which scRNA Seq data was generated included representatives of all the functional subsets identified by the flow cytometry data (Figure S1). The largest subset of Env-specific CD4+ T cells in the scRNA Seq dataset was characterized by IL-2 and CD40L protein co-expression. However, this cell subset was also a dominant population among multi-functional subsets in both the ICS and multiplexed cytokine capture assay (Figure 2D). Polyfunctional CD4+ T cells, including the subsets that correlated with reduced risk of infection in RV144 as indicated by co-expression of: IFN-γ, IL-4, IL-2, and CD154; or IL-4, IL-2, and CD154 (“correlate”), were enriched in the scRNA Seq dataset compared to the absolute frequencies by flow (Figure S1).

Using t-Distributed Stochastic Neighbor Embedding (t-SNE), we first looked to see whether dimension reduction analysis could identify gene expression clusters which corresponded to degree of functionality or protein expression of specific cytokines identified by the multiplexed cytokine capture assay (Figure 3). We overlayed the functional markers that were detected by flow cytometry on the cells clustered by gene expression. First, we looked by degree of polyfunctionality (or simply the number of functional markers co-expressed by each cell). The clusters defined by gene expression did not correspond to degree of polyfunctionality, except for a small distinct island at the top that was predominantly monofunctional. Then, we overlayed the protein expression of each cytokine/functional marker individually and found that once again they did not cluster by gene expression (Figure 3B-3E). The only exception being the one monofunctional island of cells that appeared to express only IFN-γ (Figure 3D). Lastly, we assessed whether cells with “correlate” functional cytokine profiles would cluster together by gene expression, and we found that they did not (Figure 3F).

**Figure 3.**
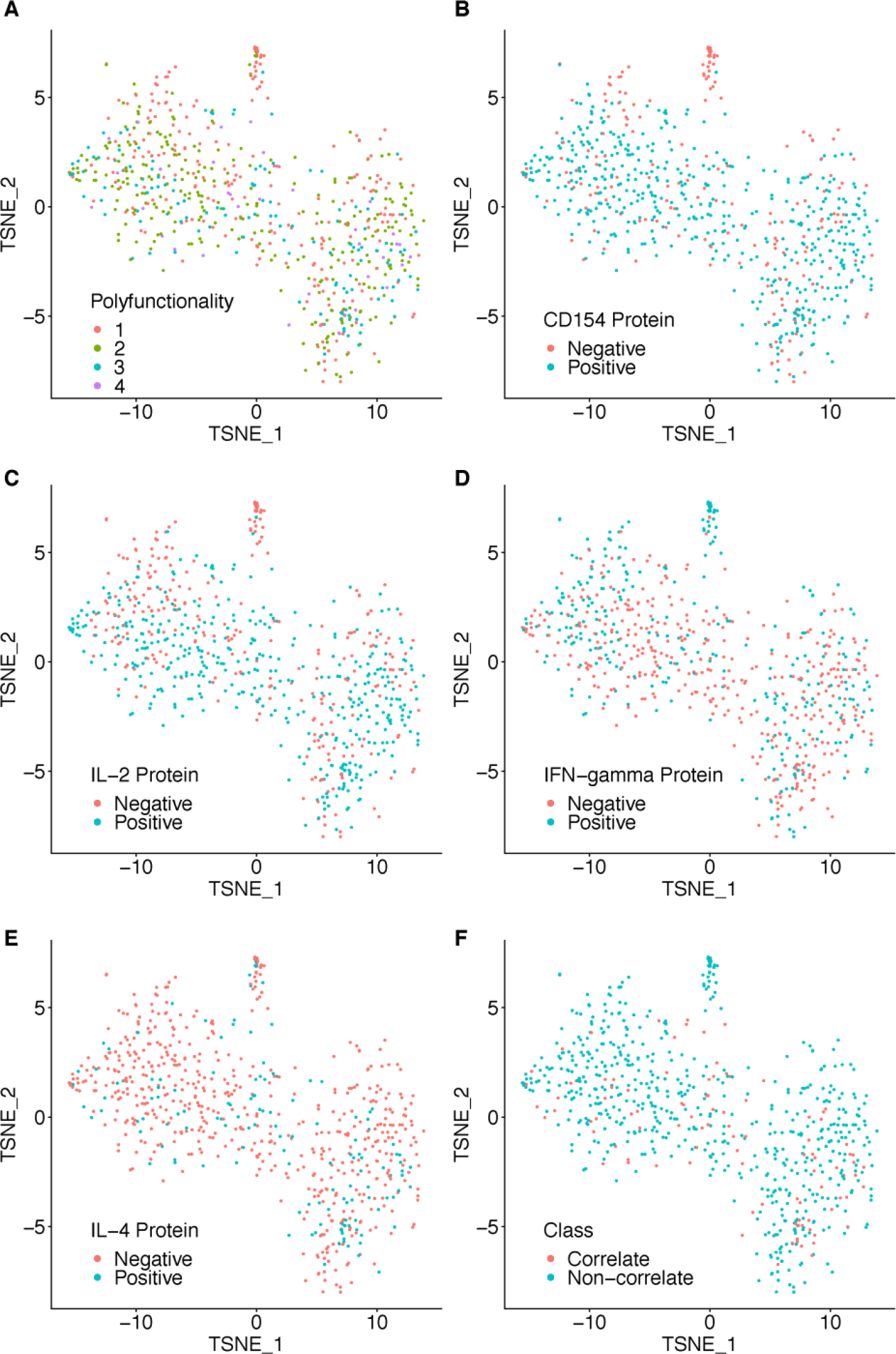
Clustering analysis of Env-specific CD4 T cells by single cell gene expression using t-Distributed Stochastic Neighbor Embedding (t-SNE). Cells are then colored by expression of cytokines or functional markers as detected by the multiplexed cytokine capture assay: A) by degree of polyfunctionality (i.e., number of contemporaneous functions/cytokines), individual functions: B) CD154, C) IL-2, D) IFN-*γ*, E) IL-4 and F) by the polyfunctional cytokine profiles associated with reduced risk of HIV infection in the RV144 vaccine trial, i.e., “correlate” versus “non-correlate”.

### Polyfunctional vaccine-specific CD4+ T cells have increased expression of genes associated with T helper function

Thus, we looked at differential gene expression analysis using MAST to determine whether there were specific genes whose expression correlated with polyfunctionality. MAST is a generalized linear framework that accounts for the bimodality of scRNA-seq data and is suitable for identifying differentially expressed genes (12). MAST models cellular detection rate as a covariate and can control for differences in abundance due to extrinsic biological and technical effects (12). We compared the monofunctional vs. polyfunctional (i.e., > 1 cytokine/function) vaccine-specific CD4+ T cells and discovered 23 genes that were significantly differentially expressed. Interestingly, the top hits included a number of protein-coding genes upregulated among the polyfunctional T cells and directly linked to CD4+ T cell function: *CD44, CD82, IFN-γ, IL-2, IL-3, IL-4, IL-5*, and *IL-21* (Figure 4A and 4B). CD44 and CD82 were interesting because they are both cell surface receptors that are upregulated by T cell activation and may be useful for identifying vaccine-induced polyfunctional T cells. CD44 plays a role in cell adhesion and cell-cell interaction and CD82 associates with CD4 and delivers costimulatory signals for the TCR/CD3 pathway. *IFN-γ, IL-2, IL-3, IL-4, IL-5*, and *IL-21* are cytokines with anti-viral or B cell activation, growth, and differentiation activity. Other transcripts that were significantly associated with polyfunctionality included multiple RNA- or protein-coding genes involved in regulation of translation (*DDX21, RPL36AL and RPL6*), secretory pathway (*CCT8*), and cell cycle/proliferation (*CYTOR, HSP90AB1*, and *MIR155HG*) (Figure 4A and 4B).

**Figure 4.**
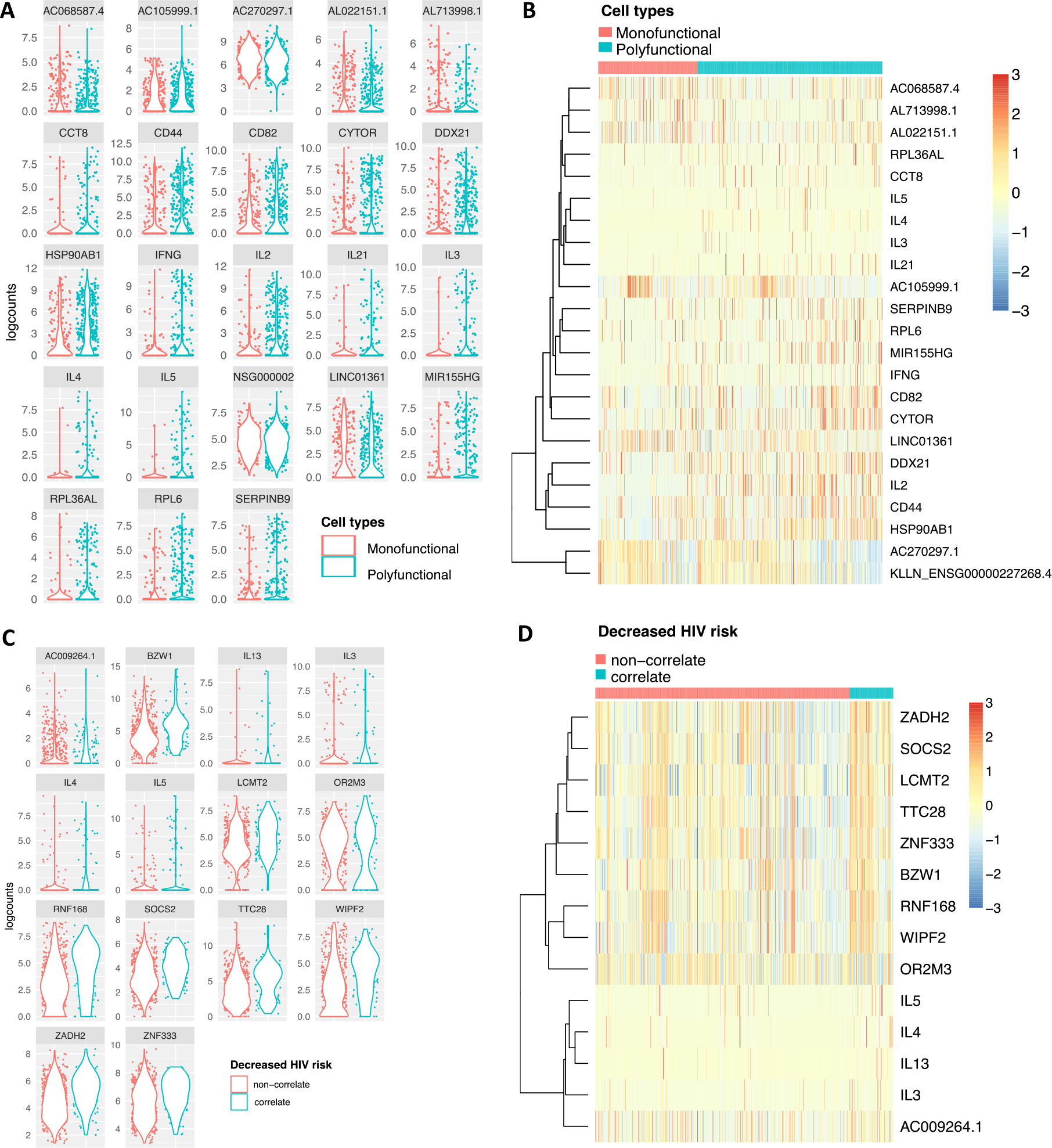
Individual genes that are differentially expressed by polyfunctional Env-specific CD4 T cells. A) Violin plots of logcounts (log2-transformed normalized counts plus a pseudo-count of 1) of the genes that were significantly differentially expressed between monofunctional (producing only 1 cytokine/function) and polyfunctional (producing 2 or more cytokines/functions) Env-specific CD4+ T cells. B) Heatmap of z-scores (mean row-centered log2-counts per million) of the same differentially expressed genes the clustered by polyfunctionality profile. C) Violin plots of logcounts of the scRNA sequences for the 14 significantly differentially expressed genes grouped by the “correlate” or “non-correlate” functional cytokine profiles among the Env-specific CD4+ T cells. D) Heatmap of z-scores of the differentially expressed genes the clustered by T cells with “correlate” or “non-correlate” cytokine profile. Genes with false discovery rate (FDR) < 0.05 and fold change of >1.25 or < -1.25 were considered significantly differentially expressed between groups.

### Th2-biased transcriptional signature predicts vaccine-specific CD4+ T cell profile correlated with reduced risk of HIV infection

Next, we compared the gene expression profiles of the cell subsets that had been previously associated with reduced risk of HIV infection in RV144 (“correlate”) compared to all other subsets of cytokine-expressing Env-specific CD4+ T cells (“non-correlate”) using MAST. We found 14 genes that were significantly differentially expressed among the Env-specific CD4+ T cells with the “correlate” compared to the “non-correlate” functional profiles. In particular, we found that expression of *IL-3, IL-4, IL-5* and *IL-13* were significantly upregulated among the T cells with the “correlate” functional profile (Figure 4C and 4D). Other transcripts that were significantly associated with the CD4+ T cells that exhibited the “correlate” functional profiles included protein-coding genes involved in regulation of JAK/STAT cell signaling pathway (*SOCS2*), transcription and/or translation (*BZW1, LCMT2 and ZNF33*), and cell cycle/proliferation (*TTC28, WIPF2)* (Figure 4A and 4B).

Building on this analysis, we asked whether there was a transcriptional signature which could predict the presence of the “correlate” functional profiles, using non-linear boosting trees implemented in XGBoost (13). To avoid overfitting and more accurately evaluate model prediction performance, we conducted 5-fold cross-validation where the data was first split into 5 smaller sets. Subsequently, each unique set was retained as the test data set, and a model was trained using the remaining 4 sets as training data and evaluated on the test set. The average Area Under the Receiver Operating Characteristic Curve (ROC AUC) for the model was 0.76 with a standard deviation of 0.10 (Figure 5A). We found that expression of *IL-4, IL-5*, and *IL-13* were among the top 30 genes which contributed to the prediction of the T cells with the “correlate” profiles (Figure 5B). We further calculated the SHAP (SHapley Additive exPlanations) values of the top 30 genes for every cell averaged across the five models to see which genes contributed to increased/decreased risk of infection (Figure 5C). SHAP values indicate a gene’s impact on a change in the model output (14). As shown in Figure 5C, high expression of *IL-5, IL-4*, and to less extent, *IL-13*, drove the model predictions towards the functional profiles correlated with reduced risk of HIV infection. These data suggest that the expression of Th2 cytokines may be uniquely detected in these polyfunctional T cell subsets and that this mechanism may be directly or indirectly associated with an immune-mediated reduced risk of infection observed in RV144.

**Figure 5.**
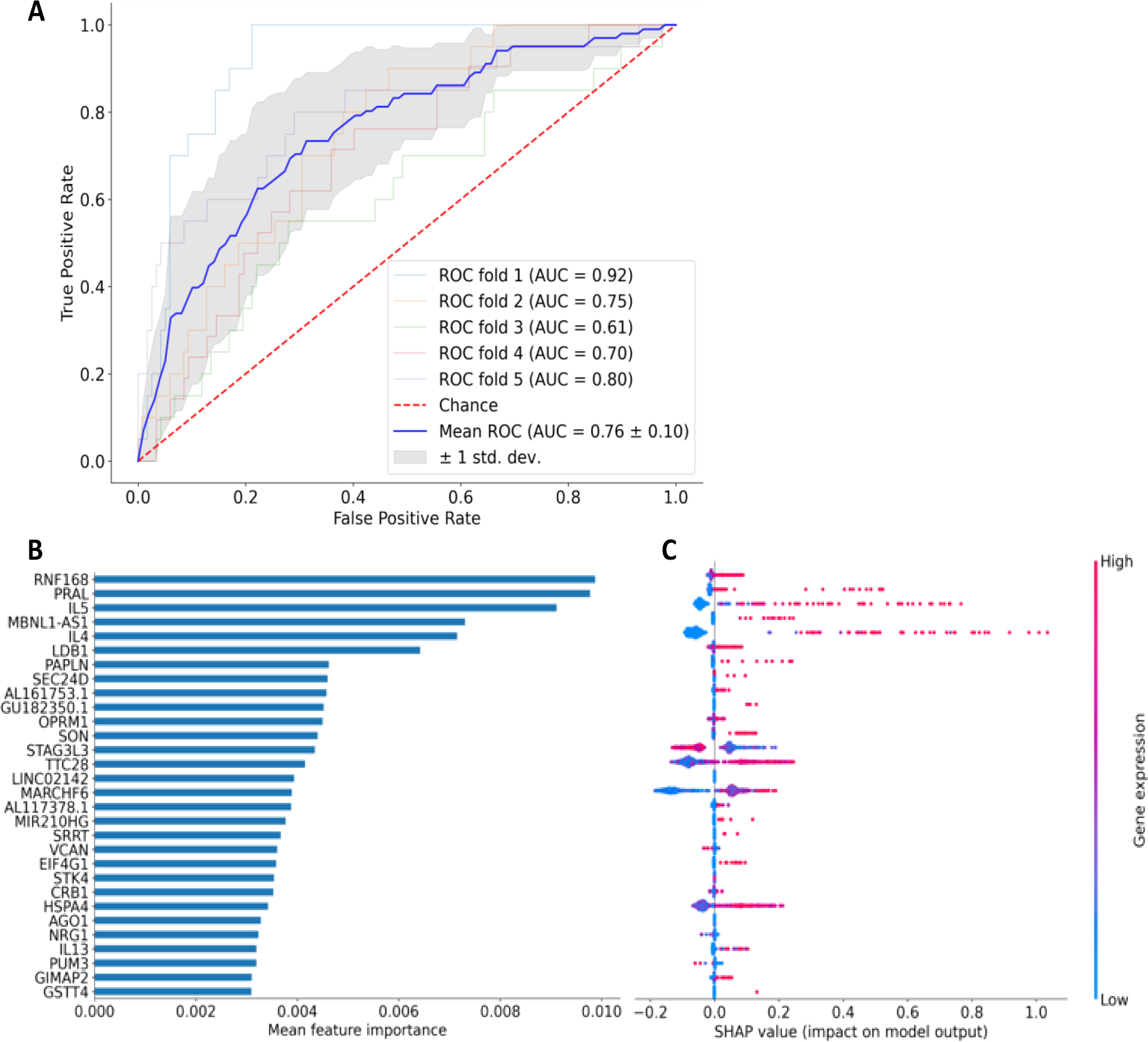
Transcriptional signatures predicted rare Env-specific polyfunctional CD4 T cell subsets that had correlated with reduced risk of HIV infection in the RV144 trial. XGBoost machine learning model was used to determine the genes that best predicted the two cytokine profiles associated with reduced risk of infection in Rv144 (“correlate” profile) versus other Env-specific CD4+ T cell subsets (“non-correlate” profile). A) Plot shows Receiver Operating Characteristic (ROC) curves created from 5-fold cross-validation. The blue line depicts the mean across the five models. B) Bar graph of the top 30 most important features based on mean feature importance across the five models. C) SHAP values showing the contribution of each gene within each cell for the top 30 genes. Color indicates the level of gene expression in each cell for the corresponding gene.

## Discussion

Thus far, the only vaccine trial to show at least partial efficacy at preventing HIV infection was RV144, in which two highly polyfunctional Env-specific CD4+ T cell subsets correlated with reduced risk of HIV infection (4, 5). Using PBMC samples from participants in the HVTN 097 clinical trial, who received the same vaccine regimen as RV144, we live-cell sorted Env-specific CD4+ T cells using a novel multiplexed cytokine capture assay and conducted scRNA sequencing to interrogate potential mechanisms underlying how Env-specific polyfunctional CD4+ T cells may have contributed to effective vaccine elicited immunity. Importantly, we found that expression of Th2 cytokines was upregulated among vaccine-specific polyfunctional T cells and specifically, in the CD4+ T cells that correlated with reduced risk of HIV infection in RV144. We also identified expression of the surface markers, *CD44* and *CD82*, as being significantly upregulated among polyfunctional Env-specific CD4+ T cells. This study represents the first time that individual polyfunctional vaccine-specific CD4+ T cells have been directly linked to their transcriptional profiles *ex vivo* and highlights potential mechanisms associated with different functional subsets of vaccine-specific CD4+ T cells.

The identification of the cell surface receptors, CD44 and CD82, as being preferentially enriched among polyfunctional CD4+ T cells suggests that they may serve as useful markers of polyfunctional T cells in future applications. In mice, CD44 is a canonical effector memory marker upregulated in response to antigen-specific activation and then maintained (15, 16). In humans, it is clear that CD44 modulates migration and extravasation and is upregulated by T cell activation or exposure to inflammatory mediators (17). Similarly, CD82 is associated with adhesion and T cell activation (18). Thus, individually or in concert, CD44 and CD82 may be particularly useful for identifying vaccine-specific polyfunctional CD4+ T cells *ex vivo*. Future studies should include evaluation of surface expression of these proteins in the context of ICS assays of vaccine-specific human CD4+ T cells to verify their ability to identify polyfunctional subsets.

Additionally, in comparing the gene expression profiles of monofunctional to polyfunctional Env-specific CD4+ T cells, we also identified an enrichment of genes that correspond to a number of cytokines: *IFN-γ, IL-2, IL-3, IL-4, IL-5*, and *IL-21*. While *IFN-γ, IL-2*, and *IL-4* correspond to proteins that were used to identify the polyfunctional T cells, *IL-3, IL-5*, and *IL-21* were not part of the capture assay panel and suggest that these functions are preferentially upregulated specifically within the multifunctional CD4+ T cells. This is interesting as these cytokines have been difficult to detect *ex vivo* among primary antigen-specific CD4+ T cells. In particular, secreted IL-21 has been previously detected after extended (40 hour) *in vitro* stimulation and was associated with improved antigen-specific T helper function supporting both antigen-specific CD8+ T cells and B cells (19, 20). Similarly, the Th2 cytokines, IL-5, and IL-13, have been difficult to detect by flow cytometry. However, the detection of *IL-5* and *IL-13* transcripts in the Env-specific polyfunctional CD4+ T cells further suggest that these T cells have enhanced capacity to help generation of effective antibody responses. This hypothesis was further supported when we interrogated the specific polyfunctional CD4+ T cells with the that had correlated with reduced risk of infection in RV144. In these analyses, the Th2 cytokines: *IL-3, IL-4, IL-5*, and *IL-13*, were significantly upregulated in the polyfunctional CD4+ T cells whose functional profiles correlated with reduced risk of infection. Furthermore, *IL-4, IL-5* and *IL-13* also formed the backbone of a transcriptional profile which predicted Env-specific CD4+ T cells with the “correlate” functional profiles.

While this study was limited in size, it provides a proof of principle that the multiplex cytokine capture assay can be used to identify, isolate, and further characterize vaccine-specific polyfunctional CD4+ T cells. The single-cell transcriptional profiling also independently confirmed the flow cytometric detection of rare unique highly polyfunctional subsets among the vaccine-specific CD4+ T cells. The gene expression analyses yielded two cell-surface receptors, CD44 and CD82, associated with T cell activation, which may be useful for identifying polyfunctional antigen-specific CD4+ T cells by flow cytometry. Lastly, we identified a Th2-biased transcriptional profile which was specific to the subset of vaccine-specific CD4+ T cells that correlated with reduced risk of HIV infection in the RV144 vaccine trial. These results support the hypothesis that the Env-specific polyfunctional CD4+ T cells induced in RV144 contributed to the reduction in HIV infection by modulating the antibody response through T cell help.

## Methods

### HVTN 097 phase 1 clinical trial samples

HVTN 097 was a randomized, double-blind, placebo-controlled phase 1b clinical trial of the RV144 vaccine regimen and was conducted in South Africa (9). Vaccine recipients received ALVAC-HIV at month 0 and 1, followed by ALVAC-HIV plus AIDSVAX gp120B/E at months 3 and 6. Peripheral blood mononuclear cells (PBMC) were collected at month 6.5 (2 weeks post fourth/last vaccination) for immunogenicity measurements, which were published previously (9). The study was approved by all appropriate IRBs and all participants provided written consent.

### Intracellular Cytokine Staining (ICS) Assay

Flow cytometry was used to examine HIV-1-specific CD4+ T cell responses from cryopreserved PBMC as previously described (9). Responses were evaluated to pools of overlapping peptides matched to the HIV-1 vaccine vector insert antigen, 92TH023-Env, measured at two weeks after final vaccination. Positivity of the ICS responses of individual cytokines or cytokine combinations was determined by a one-sided Fisher’s exact test applied to each peptide pool-specific response versus the negative control response with a discrete Bonferroni adjustment for the multiple comparisons due to testing against multiple peptide pools. Peptide pools with adjusted p-values less than 0.00001 were considered positive. We selected samples for analysis by capture assay that had a positive response to the Env peptide pool in the ICS assay during clinical trial testing.

### Multiplexed Cytokine Capture Assay

The multiplex cytokine capture assay is similar to the ICS assay by employing vaccine-specific peptides to induce the secretion of cytokines by vaccine-specific T cells (Figure 1). The catch reagents contain antibodies specific for the cytokines of interest, which adhere to the cell via a proprietary mechanism (Miltenyi Biotec), likely an antibody specific for a surface cell marker expressed on leukocytes, such as CD45, to capture the cytokines as they are secreted by the cell. After the catch reagents have bound to the cell, the cells are diluted and incubated to allow for cytokine secretion from the activated T cells. The cytokines are captured by the catch reagents and tethered to the cell that released the cytokine. The captured cytokines can then be detected by flow cytometry using surface staining methods.

PBMC were thawed quickly in warm RPMI media containing 10% FBS, L-glutamine, penicillin-streptomycin (R10) and rested overnight. An aliquot of cells was stained in Guava ViaCount reagent and counted using a Guava easyCyte (EMD Millipore), then resuspended in R10. The cells were incubated with purified anti-CD40 monoclonal antibody (Miltenyi Biotec) for 10 minutes before being stimulated (21). CD154-PE-Cy7 antibody was added along with 1 μg/mL anti-human CD28/CD49d co-stimulatory antibody and either 2 μg/mL vaccine-matched 92TH023-Env peptide pool or with peptide diluent (DMSO) for 5 hours at 37°C 5% CO2. Cells were washed in cold wash buffer (1× PBS, 2 mM EDTA, 0.5% BSA) and then incubated with 10µL per 10^6^ cells each of IFN-γ Catch Reagent (Miltenyi Biotec), IL-2 Catch Reagent (Miltenyi Biotec), IL-4 Catch Reagent (Miltenyi Biotec) in a final volume of 100 µL per 10^6^ cells for 15 min on ice. Cells were washed with R10 media and then resuspended in a minimum of 1mL per 10^6^ cells (∼10mL) of warm R10 media and incubated rotating at 37°C 5% CO2 for 1 hour. Samples were centrifuged and then washed with cold wash buffer and stained with aqua viability dye for 10 minutes on ice. Samples were washed once with cold wash buffer. The cold buffer was decanted. Samples were stained with the antibody cocktail containing lineage and phenotyping markers as well as cytokine detection antibodies and incubated for 20 minutes on ice (**Table 1**). The samples were washed once more with cold wash buffer. Samples were resuspended in 800 µL of cold buffer and filtered immediately before sorting. Live CD14- CD19- CD56- CD8- CD4+ T lymphocytes that were either CD154+ IFN-γ+ IL-2+, CD154+ IL2+, or CD154+ were single-cell sorted with a FACSAria II cell sorter. Full functional profiles were determined post-sorting using analysis of the individual cell-level flow cytometry data (acquired by indexing).

### Single-Cell RNA Sequencing (scRNA Seq)

Samples were single-cell sorted into 96-well PCR plates containing 5 µL of DNA Suspension Buffer (Teknova) with 1% BSA (Sigma-Aldrich) per well. Plates were sealed, briefly vortexed, centrifuged at 800 × g for 1 min and immediately transferred to dry ice. Plates were stored at - 80°C for at least overnight before proceeding. The plates were thawed on ice and spun at 800 × g for 1 min to collect all liquid. We used a modified version of a previously published method to amplify the whole transcriptome (11). In brief, the reverse transcriptase that is used to make cDNA has a terminal transferase activity. Then, a “template-switch” primer incorporates a second universal priming sequence yielding cDNA with two universal priming sequences.

In brief, reverse transcription occurred in two steps. The initial step was conducted in 5 µL of the lysed cell, 2.4 µM 3’ SMART CDS Primer II A (custom primer from IDT), 2.4 mM each dNTP mix (GeneAmp) and 6.32 U RNaseOUT (Invitrogen). The samples were incubated at room temperature for 1 min, 72°C for 3 min and held on ice before beginning the second step. The next step was conducted with SuperScript II Reverse Transcriptase Kit (Invitrogen), using 7 µL of the step 1 product, 1× First Strand Buffer, 2.5 mM DTT, 110 U SuperScript II and an additional 11 U RNaseOUT, as well as 1 µM SMARTer II Oligonucleotide (custom primer from IDT) and molecular-grade water to 11 µL. The samples were incubated at 42°C for 90 min, followed by a limited cycle amplification of 10 cycles at 50°C for 2 min and 42°C for 2 min, concluding with a final extension at 72°C for 15 min.

Whole transcriptome amplification (WTA) immediately proceeded with the 11 µL of the reverse transcription product, using 1× KAPA HiFi HotStart Ready Mix (Roche) and 0.5 µM IS PCR Primer (IDT). Amplification parameters were 98°C for 3 min and 22 cycles of 98°C for 15 sec, 67°C for 20 sec and 72°C for 6 min, followed by a final extension of 72°C for 5 min. The WTA product was cleaned using AMPure XP beads (Beckman Coulter). The PCR product was incubated at room temperature for 5 min with 0.8 volumes of XP beads before being placed on a magnet. The beads were washed three times with 100 µL of 80% ethanol. After briefly drying, the beads were resuspended in 20 µL of 1× TE Buffer (Sigma Aldrich) and 18 µL of supernatant was transferred to a new 96-well PCR plate. Quality and quantitation were measured by TapeStation using a High Sensitivity D5000 ScreenTape (Agilent).

Samples with a concentration ≥200 pg/µL were carried forward for sequencing according to the Nextera XT DNA Library Preparation Kit (Illumina) protocol. Samples were dual-indexed using the Nextera Index XT Kit (Illumina) and a scheme that would allow for eight pools without overlapping indices. After pooling, the concentration of each pool was measured by Qubit dsDNA HS Assay Kit (Invitrogen) and sizing was determined by Fragment Analyzer HS Small Fragment Kit (Agilent). A smear analysis was conducted in the range of 150 to 1000 bp, and the average peak size along with the Qubit results were used to determine the molarity. Each pool was then diluted to a concentration of 2 nM. The samples were sequenced on the Illumina HiSeq 2500 using PE50 on a high-output 8-lane flow cell with one pool per lane.

<u>Custom Primer Sequences, 5’ to 3’</u>

3’ SMART CDS Primer II A [Modified Oligo(dT)]:

AAGCAGTGGTATCAACGCAGAGTACT(30)VN

SMARTer II Oligonucleotide [Template Switching Oligo]:

AAGCAGTGGTATCAACGCAGAGTACATrGrGrG

IS PCR Primer [WTA PCR Primer]: AAGCAGTGGTATCAACGCAGAGT

### Statistical Methods

#### scRNA seq analysis

Fastq files were processed with Kallisto v0.45.0 (22), and gene-level counts were generated from abundances scaled using the average transcript length, averaged over samples and to library size (lengthScaledTPM), using the tximport package v1.16.1 (23). Subsequent analysis was performed using a typical Bioconductor workflow (24). Briefly, the count matrix was imported of into R to create a SingleCellExperiment object. Cells with a library size of < 30,000 reads were removed from the analysis. After QC metric filtering, a total of 689 cells were kept for downstream analysis. The logNormCounts function from the scater package v1.16.2 (25) was used to generate log-transformed normalized expression values. PCA was performed using the top 4000 highly variable genes, and tSNE (26) and UMAP (27) was subsequently performed using the top 30 principal components for data visualization. The top 30 principal components were also used to build a shared nearest-neighbor (SNN) graph, and cell clusters were identified using the walktrap algorithm from the igraph package v1.2.5 (28). Differentially expressed genes between groups were identified using MAST v1.14.0 (12). Genes with a false discovery rate (FDR) of < 0.05 and a fold change of > 1.25 or < -1.25 were considered significantly differentially expressed between groups.

### Predicting cell-level functional phenotypes correlated with decreased HIV risk

We use tree boosting, as implemented in XGBoost (13), to build a non-linear model predicting polyfunctionality from gene expression profiles. The log-transformed normalized expression values of 28,383 genes that were expressed in at least 3 cells were used to train a binary XGBoost classifier using the xgboost Python package v1.1.0 (13) with the following parameters: ‘objective’: ‘multi:softprob’, ‘num_class’: 2, ‘n_estimators’:1000, ‘n_jobs’: 6, ‘random_state’: 0, ‘scale_pos_weight’: 5.51, ‘subsample’: 0.9. The classifier was evaluated using stratified five-fold cross-validation and the Area Under the Receiver Operating Characteristic Curve (ROC AUC) metric. SHAP (Shapley Additive exPlanations) values were calculated using the shap Python v0.37.0 (14).

## Data Availability

The accession number for the RNA-sequencing data reported in this paper is Gene Expression Omnibus (GEO): GSE166945.

## Author contributions

R.G., S.D., and K.C conceived the study. K.C., C.T, and A.S. planned and developed the methods, conducted the assays, and analyzed the data. A.S Y.T. and R.A. conducted formal statistical analyses. K.C., Y.T., C.T. and A.S. drafted the original manuscript; R.G., S.D., M.J.M., and K.C. edited the manuscript. M.J.M., R.G., K.C. and S.D. secured funds and supervised the project. All authors read and approved the manuscript.

## Acknowledgements

Research reported in this publication was supported by the National Institute of Allergy and Infectious Disease of the National Institutes of Health under award numbers U19AI128914 (RG and MJM) and UM1AI068618 (MJM), as well as through VIDD Initiative awards from the Fred Hutchinson Cancer Research Center (KWC, SCD, and RG). The content is solely the responsibility of the authors and does not necessarily represent the official views of the funders. We thank the participants for volunteering their time and effort to participate in this study. We thank the HVTN 097 protocol leadership team, including Drs. Glenda Gray, Surita Roux, Nicole Grunenberg, Ying Huang, Edith Swann, and Georgia Tomaras for oversight of the clinical trial. We thank the HVTN 097 trial site staff led by Drs. Linda-Gail Bekker, Lulu Nair, Gavin Churchyard, Craig Innes, and Fatima Laher for trial implementation and collecting the samples used in this study. At the Fred Hutchinson Cancer Research Center, we thank Dr. Laura Richert-Spuhler and Stephen Voght for their assistance with the manuscript preparation.

## Tables

**Table S1.** Flow cytometry panel and cytokine capture reagents.

| Antibody target and<br>fluorochrome | Manufacturer | Clone | Catalog<br>Number | Titer (μL) |
| --- | --- | --- | --- | --- |
| Aqua Viability Dye | Invitrogen | N/A | L34957 | 0.2 |
| CD154 PE-Cy7 | BioLegend | 24-31 | 310832 | 0.15 |
| CD4 BV421 | BD Bioscience | SK3 | 566907 | 0.5 |
| CD8 Q705 | Invitrogen | 3B5 | Q10059 | 0.5 |
| CD14 BV510 | BioLegend | M5E2 | 301842 | 1.0 |
| CD19 BV510 | BD Bioscience | SJ25C1 | 562947 | 1.0 |
| CD56 APC-Cy7 | BioLegend | HCD56 | 318332 | 0.25 |
| CD45RA BV605 | BioLegend | HI100 | 304133 | 0.25 |
| CCR7 BV786 | BioLegend | G043H7 | 353226 | 2.0 |
| IL-2 APC | Miltenyi Biotec | N/A | 130-090-763 | 1.0 |
| IFN-γ FITC | Miltenyi Biotec | N/A | 130-090-433 | 1.0 |
| IL-4 PE | Miltenyi Biotec | N/A | 130-054-102 | 5.0 |

**Figure S1.**
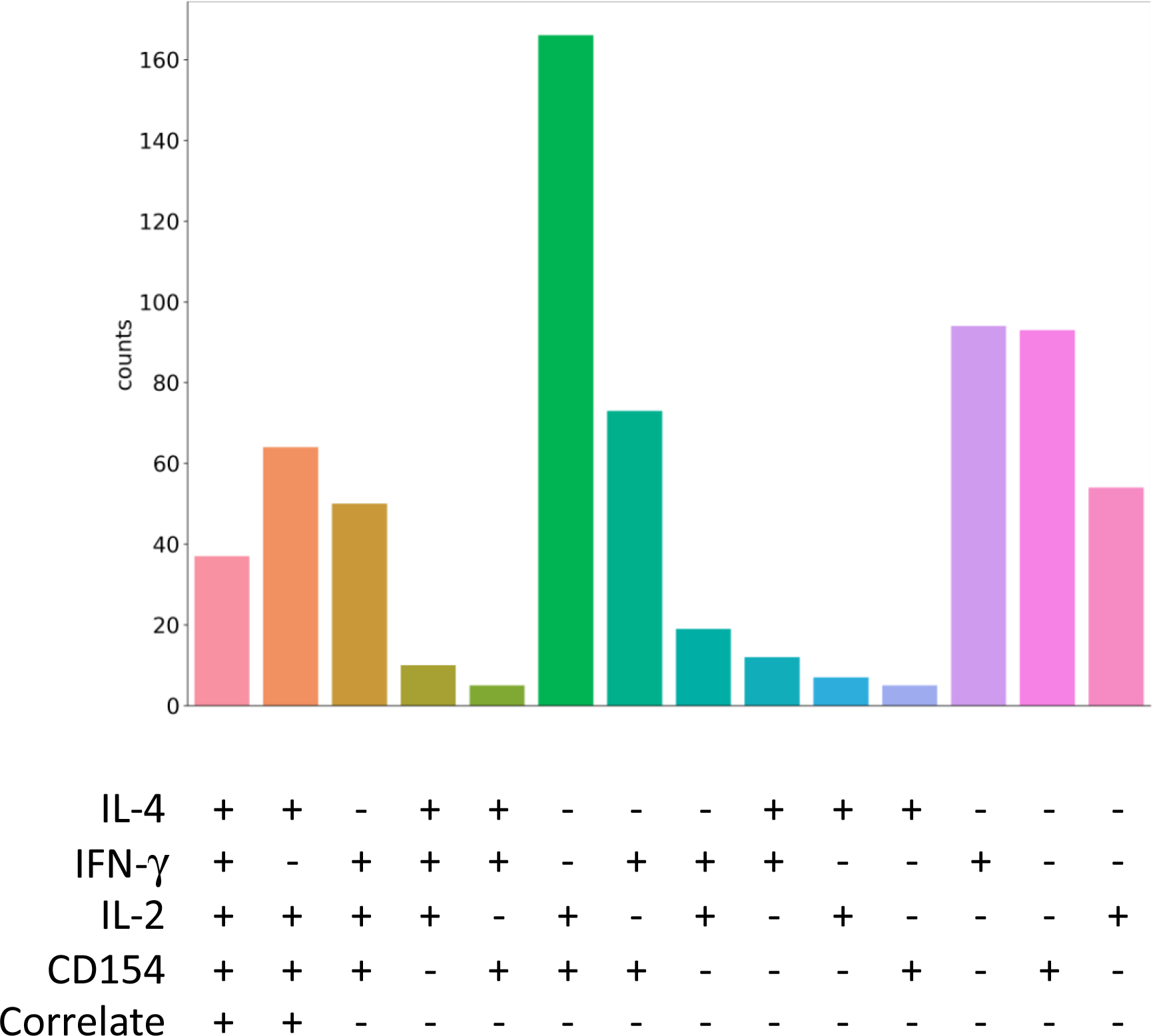
The number (count) of recovered individual Env-specific CD4+ T cells in the scRNA Seq dataset as delineated by their respective cytokine/functional protein expression determined from the multiplexed cytokine capture assay.

## Notes

### Competing Interest Statement

The authors have declared no competing interest.

